# CDS-BART: A BART-Based Foundation Model for mRNA Sequence Analysis

**DOI:** 10.64898/2026.03.09.710670

**Authors:** Erkhembayar Jadamba, Sang-Heon Lee, Jinhee Hong, Hyekyoung Lee, Sungho Lee, Hyunjin Shin

## Abstract

Summary: Recent advancements in artificial intelligence (AI) have led to the development of foundation models that interpret mRNA as a language. Notable examples include CodonBERT, hydraRNA, EVO2, and Helix-mRNA. These models demonstrate significant potential as powerful tools for mRNA research. However, to best of our knowledge, there is currently no publicly available AI model that is both easy to use and capable of analyzing mRNA sequences up to about 4kb, a length scale typical of many therapeutic mRNAs, including those encapsulated within lipid nanoparticls (LNPs). Thus, we propose CDS-BART, a user-friendly, open-source tool that integrates SentencePiece sub-word tokenization with the denoising sequence-to-sequence training of Bidirectional and Auto-Regressive Transformers (BART). CDS-BART was pre-trained on mRNA data from nine taxonomic groups provided by the NCBI RefSeq database. This comprehensive pre-training, coupled with BART’s denoising capability, ensures effective learning of codon usage, mRNA structure, evolution, and regulation. Thus, CDS-BART can ultimately deliver robust performance across a wide range of mRNA prediction tasks.

**Availability and Implementation:** CDS-BART is released under the MIT License. Latest code is available via Github at https://github.com/mogam-ai/CDS-BART.

## Introduction

Over the last few years, mRNA-based vaccines and therapeutics have become a major focus in new drug development (Pardi and Krammer, 2024). Traditionally, bioinformatics-based algorithms like RNAFold (Lorenz, et al., 2011) or LinearPartition (Zhang, et al., 2020) have been used for RNA research; however, these methods are only feasible for a limited range of downstream tasks such as mRNA secondary structure and property predictions. To overcome this limitation, large language models (LLMs) including Bidirectional Encoder Representations from Transformers (BERT), Bidirectional and Auto-Regressive Transformers (BART), and Generative Pre-trained Transformers (GPT) have inspired an approach that handles mRNA as a linguistic system with complex patterns and contextual rules. These language-based AI models serve as foundation models that enable researchers to conduct a wide range of tasks simultaneously, such as mRNA structure prediction and sequence generation (Brixi, et al., 2025; Li, et al., 2025a; Li, et al., 2024).

These mRNA language models exhibit distinctive strengths but also have limitations. For example, depending on their architectures, some LLMs struggle to process long mRNA sequences efficiently. Human mRNA coding sequences (CDS) averge around 2 kb in length, with only a small fraction likely less than 6 % exceeding 4 kb (Lopes, et al., 2021). Non-replicating mRNA vaccines encode only the target antigen and are about 2-3kb long (Zhang, et al., 2025). Even the widely used COVID-19 vaccines BNT162b2 and mRNA-1273 deliver lipid-nanoparticle(LNP)-encapsulated mRNAs of roughly 4 kb (Silva-Pilipich, et al., 2024). Similarly, gene-editing therapeutics provide a comparable benchmark, and the Streptococcus pyogenes Cas9 CDS is about 4.1 kb (Xiong, et al., 2016). Therefore, a practical 4kb upper bound covers most mRNAs used in vaccines and genome-editing..

While BERT’s masked language modeling facilitates fine-tuning for numerous downstream tasks, it is less suited for direct sequence-to-sequence (seq2seq) transformations and the analysis of very long input sequences (Devlin, et al., 2019). CodonBERT attempts to address this by tokenizing nucleotide triplets (Li, et al., 2024), thus extending the input sequence length limit. However, transcripts longer than approximately 3kb still remain out of reach. To tackle these limitations, mRNA-LM (Li, et al., 2025b), hydraRNA (Li, et al., 2025a), Helix-mRNA (Wood, et al., 2025), and EVO2 (Brixi, et al., 2025) have been proposed more recently. Among them, mRNA-LM allows the user to conduct more comprehensive analysis on 5□ UTR, coding sequences (CDS), and 3□ UTR together through contrastive language-image pre-training (CLIP) (Li, et al., 2025b). On the other hand, hydraRNA and Helix-mRNA employ a bidirectional state-space model (SSM) with or without multi-head attention for full-length RNA coverage (Li, et al., 2025a; Li, et al., 2024; Li, et al., 2025b; Wood, et al., 2025; Yang, et al., 2024). Another approach, EVO2, uses a convolutional multi-hybrid architecture, called StripedHyena2, to merge local and global contexts efficiently (Brixi, et al., 2025). However, models relying on SSM or StripedHyena2 can be complex and computationally expensive, limiting their accessibility, developability, and ease of training.

In our study, we answered these challenges by developing CDS-BART, a new mRNA foundation model that integrates SentencePiece (Kudo and Richardson, 2018) and BART (Lewis, et al., 2019). SentencePiece is a tokenization method that can compress genomic texts into far fewer non-overlapping tokens while preserving biological motifs (Zhou, et al., 2023). As a result, CDS-BART can process mRNA sequences up to about 4kb in length as input, aligning with the upper payload limit of current LNP formulations. In addition, BART’s seq2seq learning setup and denoising pre-training allow the model to take a noisy or incomplete input sentence, encode its meaning, and then generate a reconstructed input sequence that meets the task goal, unlike BERT’s token-level fill-in approach. Furthermore, our model was trained on mRNA sequence data provided by the NCBI RefSeq database including sequences of nine taxonomic groups. This diversity of the training data ensures broad coverage of complex biological patterns. In conclusion, we strongly believe that CDS-BART will serve as a practical foundation model for effective mRNA sequence property prediction, optimization, and design, particularly in the development of mRNA-based vaccines or therapeutics.

## Methods

### Dataset for pretraining

To assemble a comprehensive training dataset for pre-training, we collected mRNA sequences from the NCBI RefSeq database (Goldfarb, et al., 2024). The NCBI RefSeq database categorizes organisms into nine taxonomic groups, including archaea, bacteria, fungi, invertebrates, plants, protozoa, vertebrate mammals, vertebrate other, and viral species. Using the genome assembly summary file, we downloaded the coding sequences (CDSs) of the representative and reference genomes belonging to the nine taxonomic groups via the NCBI FTP site. The composition of CDS sequences by taxonomic group is shown in Supplementary Figure 1. After applying stringent filtering criteria, we finally obtained about 60 million CDSs for pre-training. About the filtering details, we refer the reader to the supplementary methods. The overall dataset preprocessing workflow is illustrated in Figure 1a.

**Figure 1.**
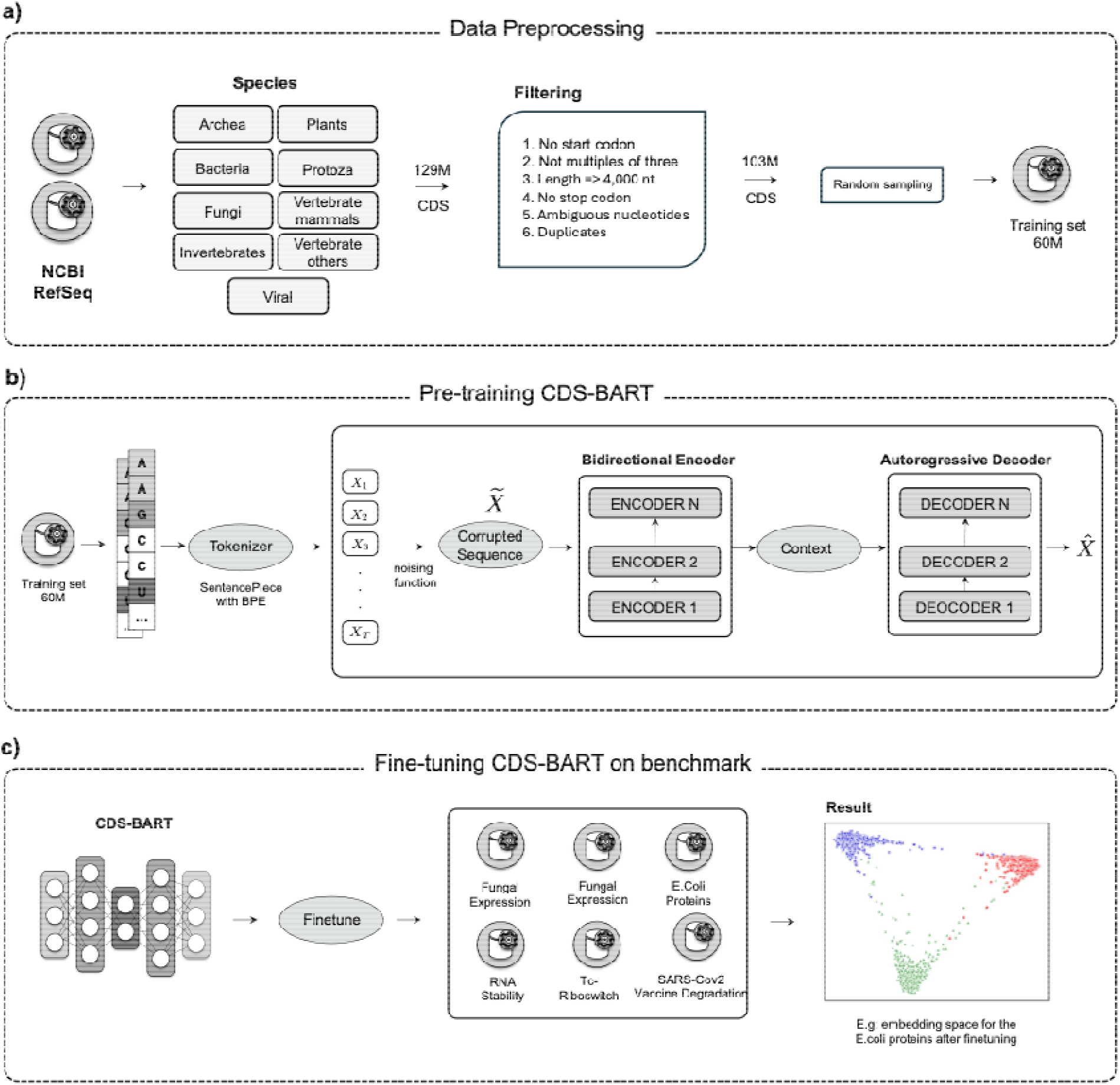
The overall procedure for establishing CDS-BART. a) In the data collection and processing step, we assembled a training dataset from the NCBI RefSeq database. This process involves the extraction of mRNA sequences from a diverse organisms categorized in nine taxonomic groups. b) CDS-BART was trained to learn mRNA sequence context by corrupting sequences with noising functions and reconstructing the original sequences. c) To evaluate the performance of CDS-BART, we fine-tuned our models on six different mRNA tasks (Supplementary Table 3). The labeled datasets were split into training, validation, and test sets in proportions of 70%, 15%, and 15%, respectively.

### Tokenizer training

To process longer sequences without increasing the model size, we developed a novel mRNA CDS tokenizer that can recognize sequence motifs of varying lengths by frequency in the training dataset. Then, we tokenized the input sequences into individual motifs using the SentencePiece method, which regards the input sequence as a continuous stream. We trained the tokenizer using byte-pair encoding (BPE) with a vocabulary size of 4,096 to process 60 million CDSs. The tokenization process required approximately 4.3 TB of memory and took two days, seven hours, and 55 minutes until completion.

### Model pretraining

CDS-BART is a BART-based denoising autoencoder built on a seq2seq architecture (Figure 1b). The model comprises two main components: a bidirectional encoder and an autoregressive decoder (Supplementary Figure 2). CDS-BART can learn the sequence context by corrupting sequences with arbitrary noising functions and reconstructing the original sequences (Supplementary Figure 3). For pre-training, we employed the BART transformer architecture, consisting of 12 encoder and decoder layers, eight attention heads, and an embedding dimension of 768. We padded or truncated CDS sequences to a maximum length of 850 tokens and constructed three different objectives: CDS-BART-seq2seq, CDS-BART-CLM, and CDS-BART-denoising. CDS-BART-seq2seq is a standard transformer encoder-decoder while CDS-BART-CLM utilizes a decoder-only architecture for next-token prediction. Finally, CDS-BART-denoising employs a masked input strategy to enhance robust contextual learning. All models were trained using the AdamW (Loshchilov and Hutter, 2017) optimizer with a learning rate of 5e-5, a weight decay of 0.01, and a linear learning rate warmup ratio of 0.1. Our model training was performed on 8 NVIDIA A100 GPUs using the HuggingFace (Wolf, et al., 2020) and DeepSpeed (Rajbhandari, et al., 2019) accelerator libraries.

### Model fine-tuning

For fine-tuning, we used six out of the seven benchmark datasets provided by CodonBERT, excluding the MLOS Flu Vaccine dataset because the metadata of this dataset did not match with the description in the CodonBERT paper. We truncated the sequences of Fungal Expression and mRNA Stability to maximum lengths of 3,000 and 1,497 nucleotides, respectively. Then, we split data according to the labeled benchmark datasets (Supplementary Table 1). Fine-tuning was conducted on a single NVIDIA A100 (80GB) GPU using the Hugging Face library and hyperparameters were also optimized based on empirical evaluations for each task (see details in Supplementary Table 2).

## Results

First, we evaluated pre-training objectives for mRNA sequence property prediction (Figure 1c), we found that the CDS-BART-denoising model significantly outperformed the other two models (CDS-BART-CLM and CDS-BART-seq2seq), achieving a Spearman correlation of 0.88 with the mRFP Expression dataset (Supplementary Figure 4). This performance underscores the effectiveness of the corruption-denoising strategy (Supplementary Figure 3), which appears to capture robust mRNA sequence patterns during training.

Next, we compared CDS-BART-denoising (hereafter referred to as CDS-BART) against other published methods, including CodonBERT (Li, et al., 2024), TextCNN (Kim, 2014), RNABERT (Akiyama and Sakakibara, 2022), RNA-FM(Chen, et al., 2022) and TF-IDF (Rajaraman and Ullman, 2011) in Table 1.

When compared to other methods, CDS-BART demonstrated notable strengths. In particular, the SARS-CoV-2 Vaccine Degradation and Tc-Riboswitch tasks are critical for understanding and improving the stability and efficacy of RNA therapeutics. In the evaluation of these tasks, CDS-BART surpassed CodonBERT, achieving remarkable improvements of 11.69% and 17.86%, respectively.

As summarized in Table 1, CDS-BART also outperformed other models in the remaining four tasks, demonstrating robust performance overall. These results show CDS-BART’s potential as a valuable tool for more accurate RNA property prediction, eventually contributing to the efficient and effective development of mRNA-based therapies and vaccines.

CDS-BART’s performance was slightly lower than CodonBERT in the Fungal Expression task (0.82 vs. 0.88). The dataset includes many fungal species with experimentally measured expression levels (Wint, et al., 2022) and shows a multi-modal label distribution (Supplementary Figure 5), reflecting strong codon-usage bias and GC-content variability among taxa. As reported in CodonBERT’s benchmarks, the representation type rather than the architecture strongly influenced performance, with codon embeddings performing better on translation-related tasks (Li, et al., 2024). In our comparative analysis, we also observed that CodonBERT’s codon-based embeddings captured fungal translation patterns slightly better. However, CDS-BART’s token-based embeddings demonstrated higher performance on more tasks, outperforming CodonBERT on five of six benchmarks.

## Discussion

In this study, we present CDS-BART, a BART-based mRNA foundation model that can extend the input sequence limit to about 4kb, which corresponds to the typical payload size for LNP delivery. CDS-BART consistently achieved high accuracy across the six benchmark tasks except for the Fungal Expression task, which is characterized by the multimodal data distribution of multiple species. The modest difference in the Fungal Expression task likely reflects representation effects, where CodonBERT’s explicit codon embeddings capture fungal-specific translation biases more directly than CDS-BART’s generalized subword embeddings. Our model also demonstrated improved results compared to CodonBERT in the SARS-CoV-2 Vaccine Degradation and Tc-Riboswitch tasks, which may require structural information for pre-training and its bidirectional-plus-autoregressive architecture. These properties are well suited to span-edited and long-context nature of therapeutic mRNA. Moreover, CDS-BART is released as an easy-to-use and openly licensed toolkit, whose pre-training and fine-tuned weights are publicly available on GitHub and Hugging Face. This will lower the barrier for researchers from basic biology to enter mRNA-vaccine engineering using AI.

On the other hand, the efficient utilization of computational resources is also critical to the development of mRNA foundation models. CDS-BART inherits the heavy computational footprint of a full encoder-decoder transformer. In detail, the dual self-attention blocks require approximately twice the memory of encoder-only model such as BERT. A single 40 GB GPU supports around 1,000 tokens, and Alibi- or RoPE-based extrapolations plateau of 2,048 tokens only at steep VRAM premium (Press, et al., 2021; Su, et al., 2021). In this work, we limited the input size to a maximum of 850 tokens obtained from only CDSs to ensure scalable processing and analysis across all datasets. Applying low-bit quantization such as QRazer and reasoning-aware distillation like DeepSeek-R1’s approach (DeepSeek-AI, et al., 2025; Lee, et al., 2025) can make CDS-BART slimer while enabling free context for longer CDSs or full RNA sequences containing CDS and UTRs together. In addition, because the same encoder-decoder backbone already powers strong text generation, a compressed CDS-BART could auto-complete missing segments or design full transcripts de novo (Supplementary Figure 6). Such capability is expected to support downstream wet-lab validation for cellular protein expression and molecular stability. Taken together, we believe that these features will position CDS-BART to become a powerful and widely adopted tool for the next wave of mRNA research, vaccine design, and therapeutic development.

