## Supplementary Figure for "CDS-BART: A BART-Based Foundation Model for mRNA Sequence Analysis"

### Supplementary Information to:

#### Supplementary Methods

1. **Data collection**

To assemble a comprehensive training dataset for pre-training, we collected mRNA sequences of a variety of organisms from the NCBI RefSeq database [1]. The assembly summary file was accessed via [NCBI Assembly Summary] (https://ftp.ncbi.nlm.nih.gov/genomes/refseq/assembly_summary_refseq.txt), and genomic sequences were downloaded from the NCBI FTP site at [NCBI Genomes] (https://ftp.ncbi.nlm.nih.gov/genomes/all/). Our data extraction process focused on obtaining all coding sequences (CDS) of representative and reference genomes. The NCBI RefSeq database categorizes organisms into nine taxonomic groups, including archaea, bacteria, fungi, invertebrates, plants, protozoa, vertebrate mammals, vertebrate other, and viral species. This classification enabled us to compile a diverse dataset comprising a total of 35,694 species, ultimately resulting in the collection of 129 million CDS (Supplementary Figure 1).

2. **Data processing**

To ensure the integrity and relevance of the mRNA sequences for subsequent analyses, we implemented a rigorous preprocessing protocol that involved stringent filtering criteria. Specifically, we excluded sequences that did not begin with a start codon (ATG), those whose lengths were not multiples of three, sequences exceeding 4 kilobases (4 kb) in length, sequences lacking stop codons (TAG, TGA, TAA), and sequences containing ambiguous nucleotides (other than A, T, G, or C). Following the application of these criteria and the removal of duplicate sequences, we finally obtained a dataset of 103 million unique CDS, which served as the foundation for further analyses. For the pretraining datasets, we randomly selected 60 million raw coding DNA sequences (CDS) for the pretraining of the CDS-BART model.

3. **Tokenizer training**

To overcome the challenge of processing longer sequences without enlarging the model, we created a novel mRNA CDS tokenizer that identifies motifs of varying lengths, informed by their frequency in the training dataset. We utilized the SentencePiece [2] method, which treats the input sequence as a continuous stream, incorporating spaces to effectively tokenize the input into individual motifs. SentencePiece operates as an unsupervised text tokenization framework that employs two subword algorithms: the Unigram language model and byte-pair encoding (BPE). The vocabulary size is established prior to program execution; hence, we trained the tokenizer using BPE with a vocabulary size of 4,096 [3]. To facilitate the training of 60 million CDS, we divided the data into chunks of 2 million sequences each. The process of training the tokenizer required approximately 4.3 TB of memory and took a total of 2 days, 7 hours, and 55 minutes to complete.

4. **Model architecture**

CDS-BART is a BART-based[4] denoising autoencoder built with a sequence-to-sequence model.

As shown in Supplementary Figure 2, CDS-BART architecture consists of two main components:

1. **Encoder**: A bidirectional encoder, like BERT, that processes the entire input sequence to understand the context of the text in both directions.

2. **Decoder**: An autoregressive decoder, like GPT [5] (Generative Pret-Training), that generates text sequentially, one token at a time, using the encoded representations.

The CDS-BART model is trained by corrupting a sequence with an arbitrary noising function and learning model to reconstruct the original sequence (Supplementary Figure 3):

1. **Corruption:** The input CDS sequences are corrupted using a noising function as masking. We deliberately masked 15% of sequences.

2. **Reconstructions**: The CDS-BART model learns to reconstruct the original, uncorrupted sequences from the noisy (masked) input.

The pretraining of CDS-BART on mRNA sequences starts with tokenizing raw sequences,

$X=(X_{1}, X_{2}, \ldots, X_{T}$)	(1)

where $X_{j}$represents individual tokens, and *T* is the sequence length. Then CDS-BART applies a **noising function**, to *X*, producing a corrupted sequence $\tilde{X}$. We used masking of input sequences as our default noising function,

$f_{noise}\left( X \right)= \tilde{X}.$(2)

The corrupted sequence $\tilde{X}$ is fed into the **bidirectional encoder**, which maps *H* to contextual embeddings:

$$H=Encoder\left( \tilde{X} \right). (3)$$

Here, $H \in\mathbb{R}^{Txd}$ , where *d* is the hidden size of the encoder, and each $h_{t}$ (a row of *H*) represents the contextualized embedding for the token $X_{t}$.

The training objective is to maximize the log likelihood of reconstructing the original sequence *X* from the corrupted input $\tilde{X}$:

$\mathcal{L=}\mathbb{E}_{X\sim\mathcal{D}} \left[ log P\left( X \right|\tilde{X}) \right]$	(4)

Where *D* is the training dataset. The probability is computed as:

$P\left( X \right|\tilde{X})= \prod_{t=1}^{T} P\left( x_{t} \right| x_{<t}, H)$	(5)

5. **Pretraining detail**

We employed a BART transformer architecture for all pretraining experiments. The base configuration consists of 12 blocks encoder and decoder layers, 8 attention heads, an embedding dimension of 768. CDS sequences were padded or truncated to a maximum length of 850 tokens.

We pretrained three models using distinct objectives.

- **CDS-BART-seq2seq (Sequence-to-sequence (Seq2Seq) autoencoder).** This model was trained as a standard transformer encoder-decoder. The input to the encoder was the complete, unprocessed CDS sequence (prepended with [<bos>] and appended with [<eos>]). The decoder was tasked with autoregressively generating the identical sequence, conditioned on the encoder's output and the previously generated tokens. The objective was to minimize the cross-entropy loss between the decoder's output logits and the target sequence.
- **CDS-BART-CLM (Causal Language Model).** For the CDS-BART-CLM, we utilized a decoder-only transformer architecture. The model was trained to predict the next nucleotide token given the preceding tokens in the sequence. The objective was to minimize the standard cross-entropy loss for next-token prediction across the entire sequence.
- **CDS-BART-denoising (Denoising Seq2Seq).** For this model we used the same transformer encoder-decoder as seq2seq. However, the input CDS fed to the encoder was corrupted using a masking strategy. Specifically, 10% of the input tokens in each sequence were randomly selected and replaced with the mask token.

All models were trained using the AdamW[6] optimizer with a learning rate of 5e-5, and weight decay 0.01. We employed a linear learning rate warmup ratio of 0.1. A batch size 16 training and validation data was used. The pre-training process was performed on 8 x NVIDIA A100 (80GB) GPUs using HuggingFace [7] and DeepSpeed [8] accelerator library.

In addition, we assessed three different pretraining objectives, CDS-BART-Denoising, CDS-BART-seq2seq, and CDS-BART-CLM, to investigate the relative advantages of these objectives in fine-tuning. The pretraining objectives were evaluated with the mRFP Expression dataset, yielding performance scores (i.e., Spearman correlation) of 0.640, 0.798, and 0.875, respectively. The results suggest that the higher performance of CDS-BART-Denoising is likely due to its corruption-based pretraining strategy, which encourages the model to learn more robust contextual representations by recovering the corrupted tokens. This form of denoising forces the encoder to capture richer and more robust bidirectional dependencies within mRNA sequences, translating well to fine-tuning scenarios. In contrast, CDS-BART-seq2seq a seq2seq autoencoder, rather focuses on simply reconstructing the input, which may offer less representational challenges and diversity for homogeneous mRNA sequences. Lastly, CDS-BART-CLM, the decoder-only CLM, although known to be effective for sequence generation based on left-to-right context, may not be optimal for such tasks as requiring a global understanding of long mRNA sequences. These findings highlight the efficacy of the denoising pretraining strategy in enhancing the model's ability to generalize across different mRNA-related tasks.

6. **Finetuning dataset**

CodonBERT provides seven types of datasets, including MLOS Flu Vaccines, mRFP Expression [9] E. coli Proteins [10], mRNA Stability [11], Tc-Riboswitches [12], Fungal Expression [13], and SARS-CoV-2 Vaccine Degradation [14], which encompass diverse biological data that form a robust foundation for analyzing gene expression and protein production across various organisms and conditions. In our study, the MLOS Flu Vaccine dataset [15] was excluded due to discrepancies in the number of data points compared to the original paper and the lack of division into training and validation sets, resulting in the utilization of six out of the seven available datasets. Furthermore, the Fungal Expression and mRNA Stability datasets contained more data points than reported in the original paper; therefore, we truncated the sequences to a maximum length of 3,000 nucleotides to ensure consistency with CodonBERT. Subsequently, we employed the labeled datasets to perform the split into training, validation, and test sets, adhering to proportions of 70%, 15%, and 15%, respectively.

7. **Finetuning detail**

In our finetuning experiments, we utilized the Hugging Face library, leveraging a pretrained BART transformer model. The finetuning process was conducted on a single NVIDIA A100 (80GB) GPU and aimed to optimize performance across five distinct tasks, as summarized in the accompanying finetuning dataset (supplementary table 1). The following hyperparameters were selected based on empirical evaluations to achieve optimal performance on the respective tasks. The detailed hyperparameters for the best-performing models across each task are shown in the Supplementary Table 2, which provides a comprehensive overview of the configurations employed for each dataset.

8. **Embedding space visualization details**

To evaluate the effectiveness of the pretrained model in capturing sequence patterns across different taxonomic groups, we encoded the data and projected it into a two-dimensional space to analyze clustering behavior relative to taxonomic classification. For this study, we selected four taxonomic groups plant, bacteria, invertebrate, and fungi from a total of nine groups, randomly sampling 500 CDS sequences from each. These sampled sequences were processed using the CDS-BART model to generate embeddings. Subsequently, we employed Principal Component Analysis (PCA) to reduce the dimensionality from 768 features to 50 features, facilitating more efficient data handling. Following this, Uniform Manifold Approximation and Projection (UMAP) was applied for projection, allowing for visualization of the clustering patterns. This approach revealed that sequences belonging to the same taxonomic group clustered together, delineating distinct boundaries, as illustrated in Supplementary Figure 6a.

#### Supplementary Figures

**Supplementary Figure 1.** NCBI raw data taxonomy distribution used for pretraining.

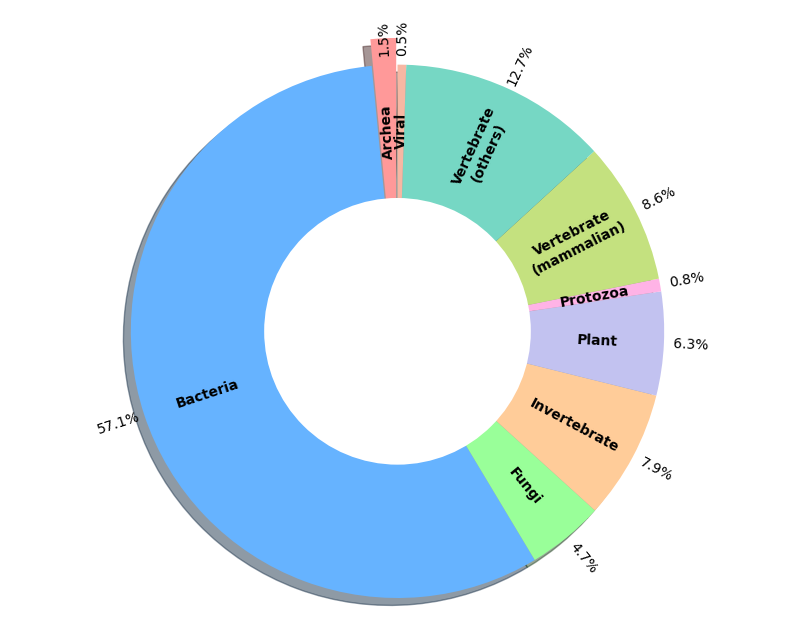

**Supplementary Figure 2**. CDS-BART bidirectional encoder and autoregressive decoder.

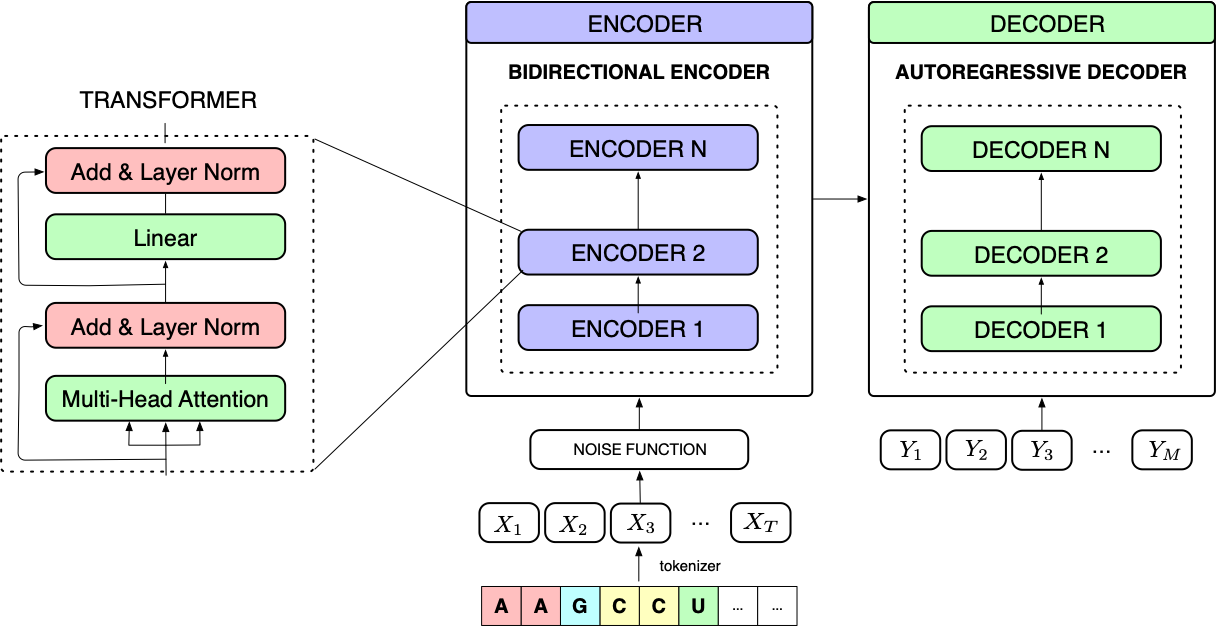

**Supplementary Figure 3**. CDS-BART pretraining process with corruption and reconstruction

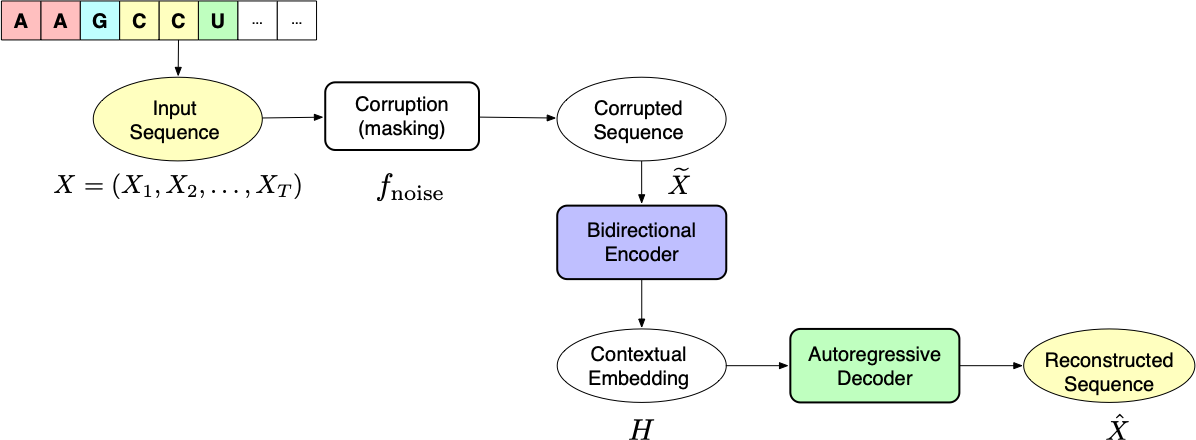

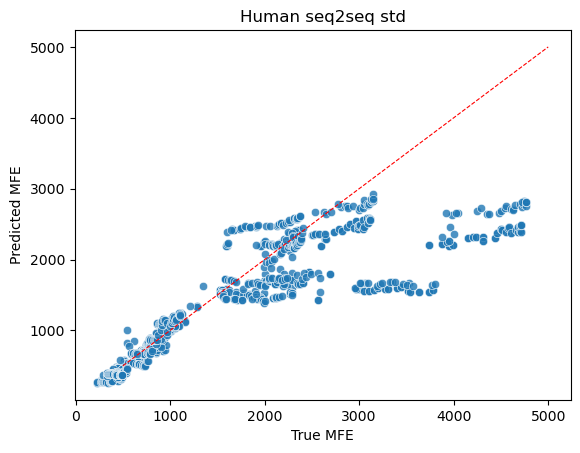
**Supplementary Figure 4.**

Comparing the results of three types of pretraining models CDS-CLM, CDS-Seq2Seq, CDS-denoise specifically focusing on their performance on the mRFP Expression dataset.

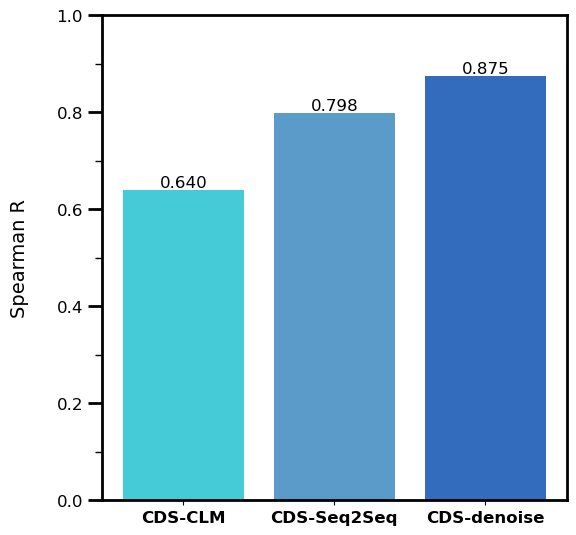

**Supplementary Figure 5.** Task-specific label distributions for prediction targets.

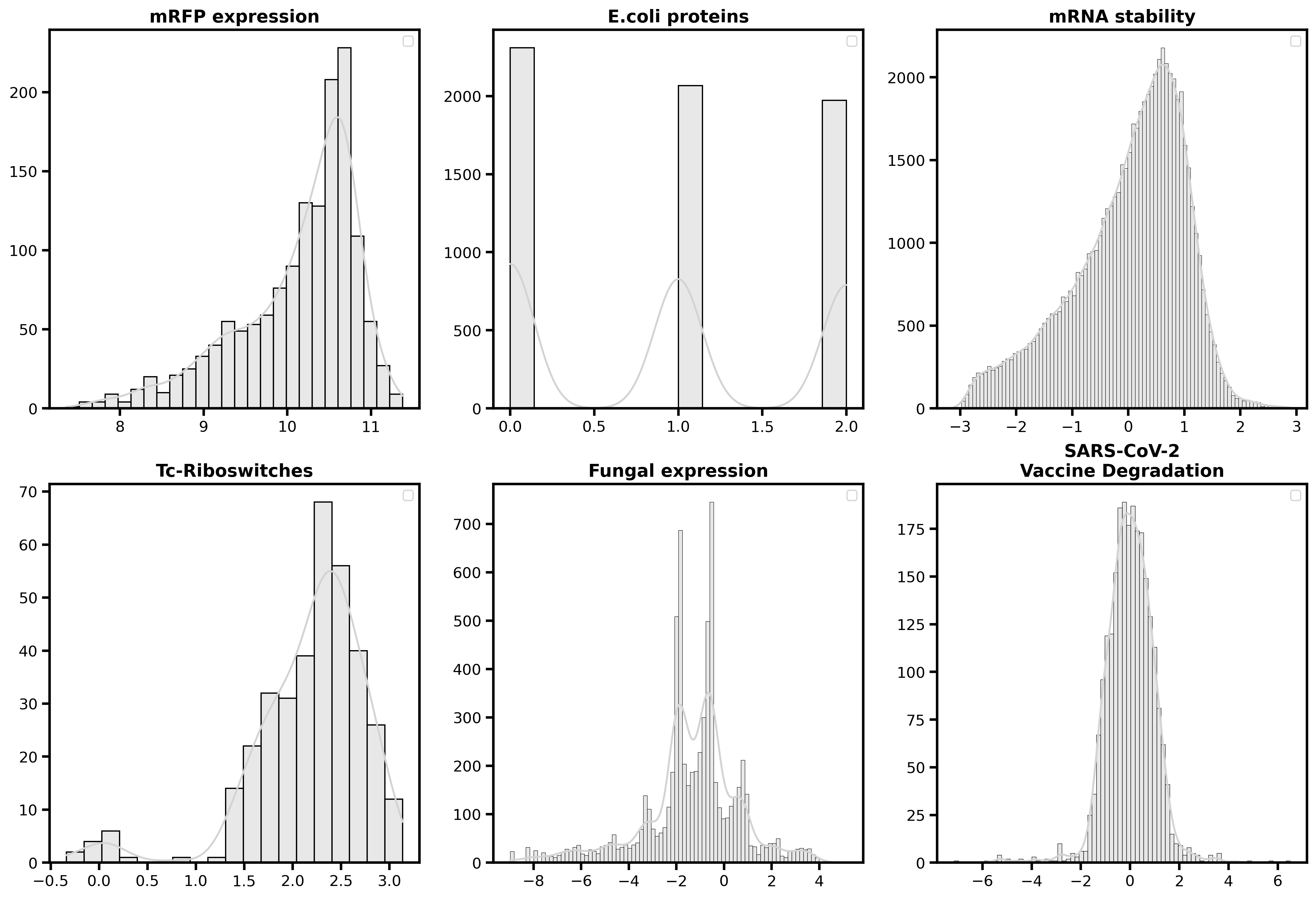

**Supplementary Figure 6.** CDS-BART embedding space representation.

a) The embedding space of the pretraining model for four NCBI taxonomies is represented using UMAP

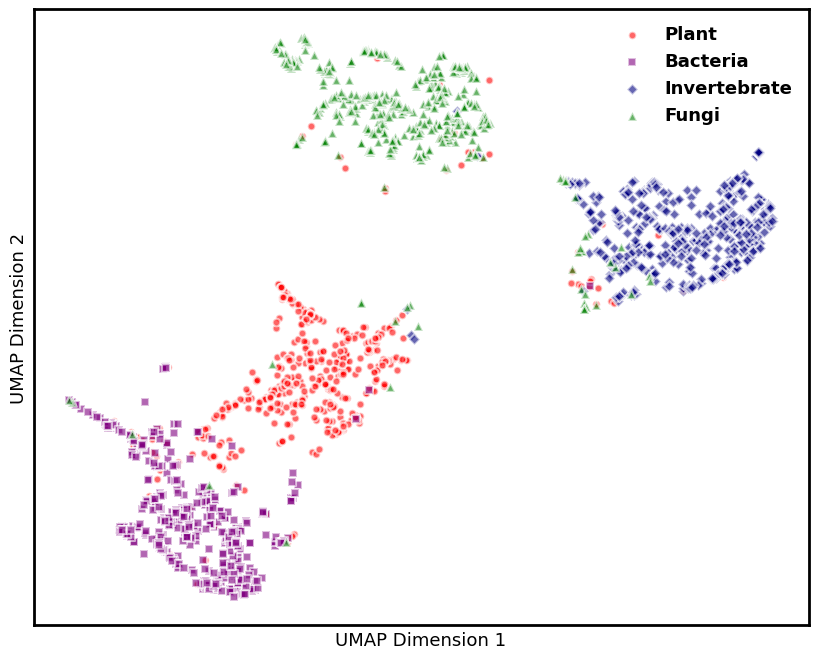

b) The finetuned model’s embedding space for the E.coli proteins classification task is represented using UMAP. E.coli proteins are categorized into low, medium and high expression levels.

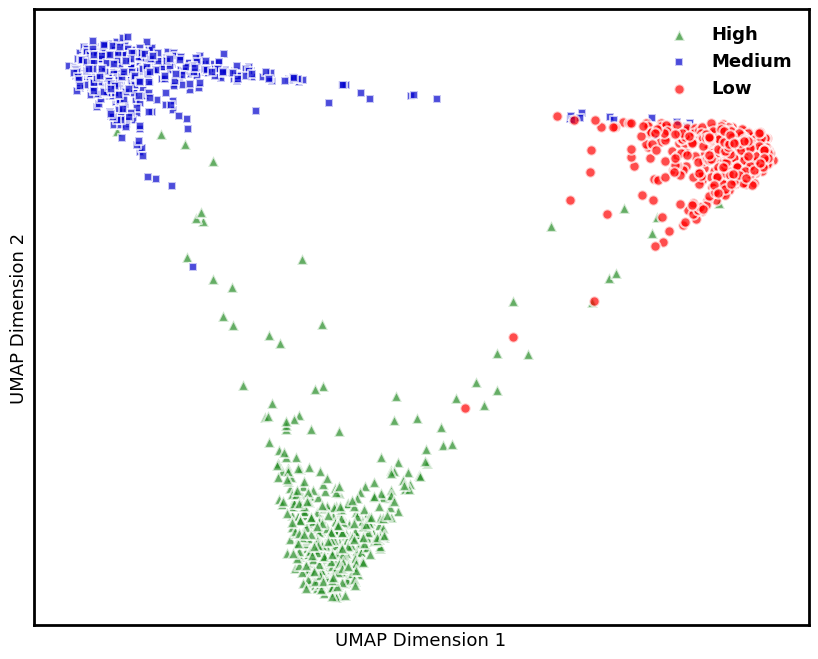

**Supplementary Figure 7:** Comparison of CDS-BART with prior methods across six benchmark tasks.

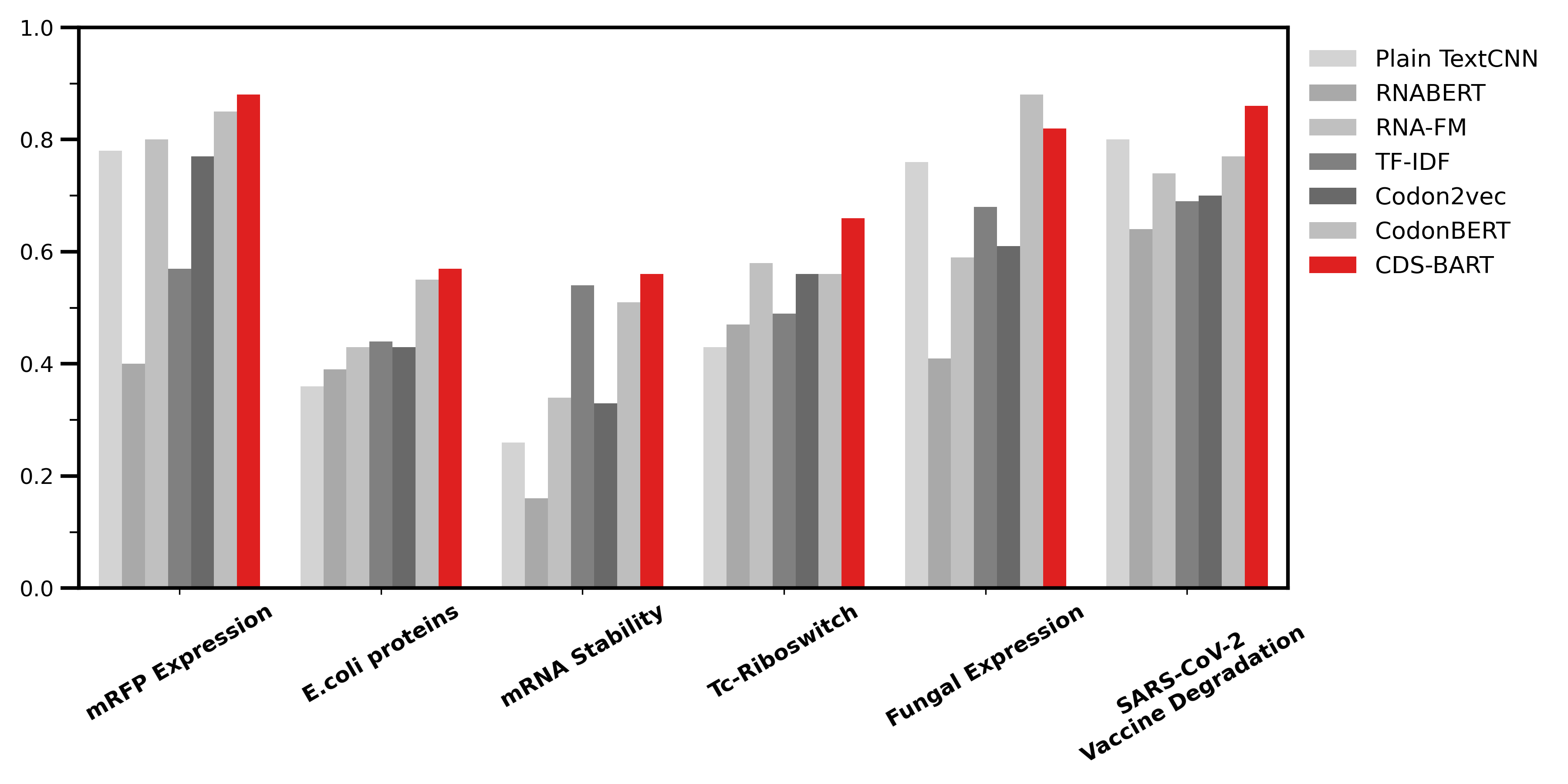

**Supplementary Figure 8.** Fine-tuning train/evaluation loss.

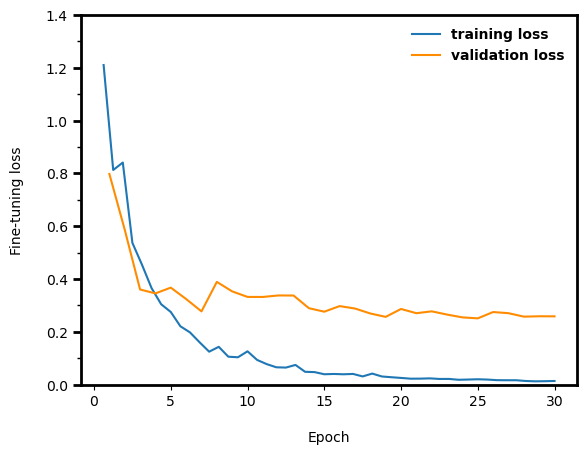

**Supplementary Figure 9.** Performance distribution of CDS-BART with five random seeds for each task. This distribution demonstrates robust and stable performance.

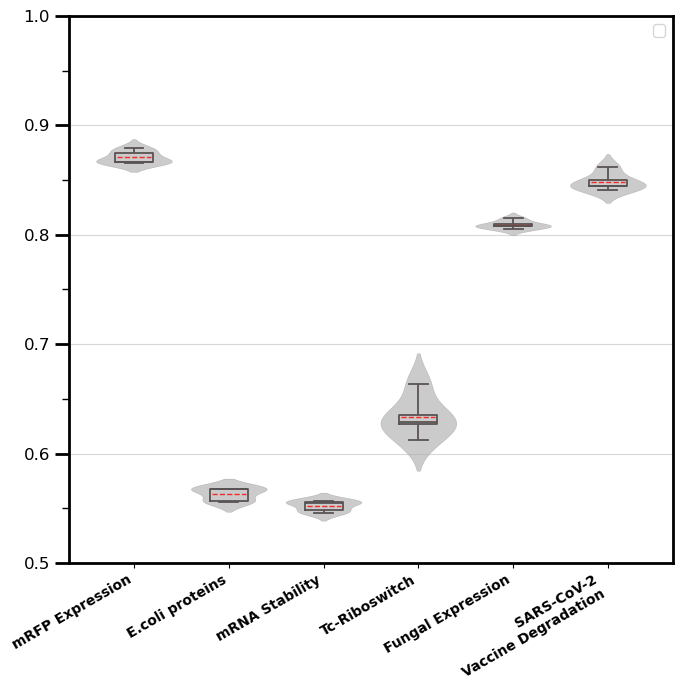

#### Supplementary Tables

**Supplementary Table 1.** The details of the benchmark dataset used in evaluation of the CDS-BART

| **Data set** | **Target** | **Category** | **No. of mRNAs** | **Seq length** |
| --- | --- | --- | --- | --- |
| mRFP expresssion [9] | Expression | Regression | 1459 | 678-678 |
| *E. coli* proteins [10] | Expression | Classification | 6,348 | 171-3000 |
| mRNA stability [11] | Stability | Regression | 41,063 | 30-1497 |
| Tc-Riboswitches [12] | Switching factor | Regression | 355 | 66-75 |
| Fungal expression [13] | Expression | Regression | 7,053 | 150-3000 |
| SARS-CoV-2 vaccine degradation [14] | Degradation | Regression | 2,400 | 81-81 |

Each data set is spilt into training, validation, and test with a 0.7, 0.15, and 0.15 ratio. All the methods were optimized on the same data split.

**Supplementary Table 2.** Optimal hyperparameter configurations across six benchmark datasets

| **Task** | **Learning rate** | **Batch size** | **Warmup ratio** | **Weight decay** | **Learning rate**  **schedular** | **Epoch** | **Normalization** |
| --- | --- | --- | --- | --- | --- | --- | --- |
| mRFP expression | 2e-5 | 4 | 0.1 | 0.01 | Linear | 30 | Standard Scaling |
| E.coli proteins | 1e-5 | 64 | 0.1 | 0.01 | Cosine | 30 | - |
| mRNA Stability | 2e-5 | 128 | 0.1 | 0.01 | Cosine | 30 | - |
| Tc-Riboswitch | 1e-5 | 4 | 0.1 | 0.01 | Linear | 30 | - |
| Fungal Expression | 2e-5 | 64 | 0.1 | 0.01 | Linear | 30 | - |
| SARS-CoV-2  Vaccine Degradation | 1e-5 | 4 | 0.1 | 0.01 | Linear | 30 | - |

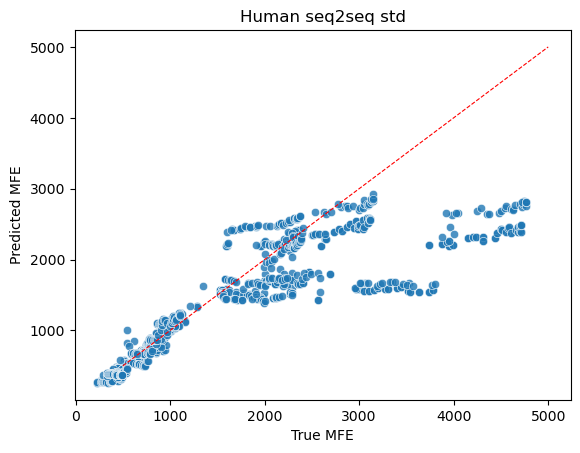
