## Supplementary material for "CDS-BART: A BART-Based Foundation Model for mRNA Sequence Analysis": Table 1

**Table 1.** Results of the evaluation on classification/regression of the benchmark datasets.^a^

| **Model** | **mRFP Expression** | **E.coli  proteins** | **mRNA  Stability** | **Tc- Riboswitch** | **Fungal Expression** | **SARS-CoV-2 Vaccine  Degradation** |
| --- | --- | --- | --- | --- | --- | --- |
| **Dataset Task** | Regression | Classification | Regression | Regression | Regression | Regression |
| Nucleotide-based |  |  |  |  |  |  |
| plain TextCNN | 0.62 | 0.39 | 0.01 | 0.41 | 0.53 | 0.55 |
| RNABERT+TextCNN | 0.4 | 0.39 | 0.16 | 0.47 | 0.41 | 0.64 |
| RNA-FM+TextCNN | 0.8 | 0.43 | 0.34 | 0.58 | 0.59 | 0.74 |
| Codon-based |  |  |  |  |  |  |
| TF-IDF | 0.57 | 0.44 | 0.54 | 0.49 | 0.68 | 0.69 |
| plain TexTCNN | 0.78 | 0.36 | 0.26 | 0.43 | 0.76 | 0.8 |
| Codon2vec+TextCNN | 0.77 | 0.43 | 0.33 | 0.56 | 0.61 | 0.7 |
| CodonBERT | 0.85 | 0.55 | 0.51 | 0.56 | **0.88** | 0.77 |
| CDS-BART | **0.88** | **0.57** | **0.56** | **0.66** | 0.82 | **0.86** |

^a^ Comparison of CDS-BART to prior methods on six downstream tasks. For regression tasks, the corresponding Spearman’s rank correlation values were used. For the classification task (E. coli proteins data set), classification accuracy was calculated. The best value of correlation and accuracy for each task was marked in bold.
